# Rab28 is localised to photoreceptor outer segments, regulates outer segment shedding but is not linked to retinal degeneration in zebrafish

**DOI:** 10.1101/720029

**Authors:** Stephen P. Carter, Ailís L. Moran, David Matallanas, Gavin J. McManus, Oliver E. Blacque, Breandán N. Kennedy

## Abstract

The photoreceptor outer segment is the canonical example of a modified and highly specialised cilium, with an expanded membrane surface area in the form of discs or lamellae for efficient light detection. Many ciliary proteins are essential for normal photoreceptor function and cilium dysfunction often results in retinal degeneration leading to impaired vision. Herein, we investigate the function and localisation of the ciliary G-protein RAB28 in zebrafish cone photoreceptors. CRISPR-Cas9 generated *rab28* mutant zebrafish display a reduction in shed outer segment material in the RPE at 1 month post fertilisation (mpf), but otherwise normal retinal structure and visual function up to 12 mpf. Cone photoreceptor-specific transgenic reporter lines show Rab28 localises almost exclusively to outer segments, independently of nucleotide binding. Co-immunoprecipitation analysis demonstrates tagged Rab28 interacts with components of the phototransduction cascade, including opsins, Phosphodiesterase 6C and Guanylate Cyclase 2D. Our data shed light on RAB28 function in cones and provide a model for RAB28-associated cone-rod dystrophy.

## Introduction

The photoreceptor outer segment (OS) is an elaborate membranous organelle which functions in the detection of light stimuli and their conversion to electrical signals via phototransduction (Sung & Chuang 2010). Outer segments are modified primary cilia and as such the molecular machinery which regulates transport and signalling within cilia is also essential for OS formation and function (Wheway et al. 2014). Furthermore, blindness due to photoreceptor degeneration (PRD) is a common phenotype of genetic diseases known as ciliopathies, characterised by ciliary dysfunction (Waters & Beales 2011; Bujakowska et al. 2017).

Photoreceptor OS are composed of flattened, closed disks surrounded by an outer membrane in rods and open lamellae in cones. New discs/lamellae form at the base of the OS as ciliary ectosomes (Ding et al. 2015; Salinas et al. 2017) and gradually migrate upwards as the oldest discs/lamellae at the OS tip are shed and phagocytosed daily by the retinal pigment epithelium (RPE). OS shedding is integral to photoreceptor health and survival: as the OS are exposed to high levels of light, the oldest discs/lamellae accumulate photo-oxidatively damaged compounds (Sung & Chuang 2010). Despite its essential role in photoreceptor biology, the molecular machinery which regulates OS shedding is poorly described. Recently, knockout of the small GTPase RAB28 resulted in impaired shedding from the tips of mouse cones, but not rods (Ying et al. 2018). Failure to shed old lamellae led to the accumulation of membranous material at cone tips and eventual degeneration and death of the cones, followed by rods. In humans, *RAB28* null and hypomorphic alleles cause autosomal recessive cone-rod dystrophy (arCRD) (Roosing et al. 2013; Riveiro-Álvarez et al. 2015; Lee et al. 2017), to our knowledge the only example of inherited PRD arising exclusively from a disorder of cone OS (COS) shedding.

In *C. elegans*, we previously demonstrated that RAB28 is an IFT and BBSome associated ciliary protein (Jensen et al. 2016). Here, we generate zebrafish *rab28* knockout and transgenic reporter models to investigate the localisation, function and interactome of RAB28 in cone photoreceptors, and the dependency of these on nucleotide binding. Localisation of RAB28 to the OS is partially dependent GTP/GDP-binding, overexpression of GTP-preferring RAB28 in cones results in visual behaviour defects and RAB28 biochemically associates with components of the phototransduction cascade, as well as vesicle trafficking and mitochondrial proteins. Furthermore, *rab28* null zebrafish display reduced OS shedding from at least 1 month post fertilisation (mpf), but not retinal degeneration.

## Materials and Methods

### Zebrafish strains and maintenance

Zebrafish larvae from 0-5 days post fertilisation (dpf) were cultured in Petri dishes of E2 medium (0.137 M NaCl, 5.4 mM KCl, 5.5 mM Na_2_HPO_4_, 0.44 mM KH_2_PO_4_, 1.3 mM CaCl_2_, 1.0 mM MgSO_4_ and 4.2 mM NaHCO_3_, conductivity ~1500 μS, pH 7.2) at 27°C on a 14 h/ 10 h light-dark cycle.

Adult zebrafish were housed in 1.4, 2.8 or 9.5 L tanks in system water and maintained at a temperature of 27°C on a 14 h/10 h light-dark cycle. The UCD facility environmental parameters are reported at (Crowley et al. n.d.). Juvenile fish were fed an increasingly complex, specialised diet (Special Diet Services) and gradually transferred to a diet of mainly brine shrimp (Artemia sp.). Zebrafish strains used in this study were: WT (Tü), *rab-28^ucd7^, rab-28^ucd8^*, Tg[gnat2:eGFP], Tg[gnat2:eGFP-rab28], Tg[gnat2:eGFP-rab28^Q72L^] and Tg[gnat2:eGFP-rab28^T26N^]

### Ethics statement

All animal experiments were conducted with the approval of the UCD Animal Research Ethics Committee (AREC-Kennedy) and the Health Products Regulatory Authority (Project authorisation AE18982/P062). All experiments were performed in accordance with relevant guidelines and regulations.

### Generation of *rab28* mutant zebrafish

sgRNAs were designed using the ZiFiT Targeter (v4.2) online tool. Several sgRNAs were designed against the zebrafish *rab28* cDNA sequence. The sgRNA against exon 2 of *rab28* was chosen as there was sufficient genomic sequence data to facilitate genotyping. sgRNAs were cloned into the pDR274 vector (Addgene) following a previously described protocol (Hwang et al. 2013). CRISPR mutants were generated by microinjection of Cas9-sgRNA ribonucleoprotein particles (RNPs) into one-cell stage WT embryos (Cas9 protein was acquired from Integrated DNA Technologies). P_0_ injected fish were raised to adulthood and screened for germline transmission of potential *rab28* null alleles. These were outcrossed to a WT line and the subsequent heterozygous F_1_ fish raised and in-crossed to generate homozygous *rab28*^-/-^ larvae.

### Zebrafish transgenesis

Transgenic zebrafish expressing eGFP-Rab28 in cone photoreceptors were generated by microinjection of plasmids containing a Tol2-gnat2:eGFP-rab28(cDNA)-Tol2 construct, together with Tol2 transposase mRNA. Plasmids were generated by MultiSite Gateway cloning using the Tol2kit and following a previously described protocol (Kwan et al. 2007). The *gnat2* promoter was cloned previously (Kennedy et al. 2007). The zebrafish *rab28* cDNA clone was acquired from the Zebrafish Gene Collection (IMAGE ID: 2643307). The T26N (GDP-preferring) and Q72L (GTP-preferring) mutants of RAB28 were generated by site-directed mutagenesis of the cDNA. Injected embryos were treated with 75 μM phenylthiourea (PTU, Sigma) diluted in embryo medium to suppress melanogenesis and screened for expression of eGFP at 5 dpf. Those larvae positive for eGFP were raised to adulthood and outcrossed to a WT line to generate heterozygous F_1_ transgenic carriers.

### Molecular biology

sgRNAs and Tol2 transposase mRNA were generated by *in vitro* transcription using the MEGAshortscript and mMessage mMachine SP6 kits (Invitrogen), respectively, following the manufacturer’s protocol. RNA was purified by LiCl precipitation. For RT-PCR, 5 dpf zebrafish larvae were placed in RNAlater and stored at 4°C overnight. Larvae were homogenised by aspiration through a needle and syringe and RNA was extracted from the resulting lysate using the mirVana RNA isolation kit (Life Technologies), following the manufacturer’s protocol. RNA was subsequently purified and concentrated by ethanol precipitation. cDNA was generated from the RNA using the RevertAid cDNA synthesis kit (ThermoFisher). This cDNA was then used as template DNA for subsequent PCR reactions. Primers used in genotyping and generating transgenic constructs are provided in **Table 1**.

**Table 1.**
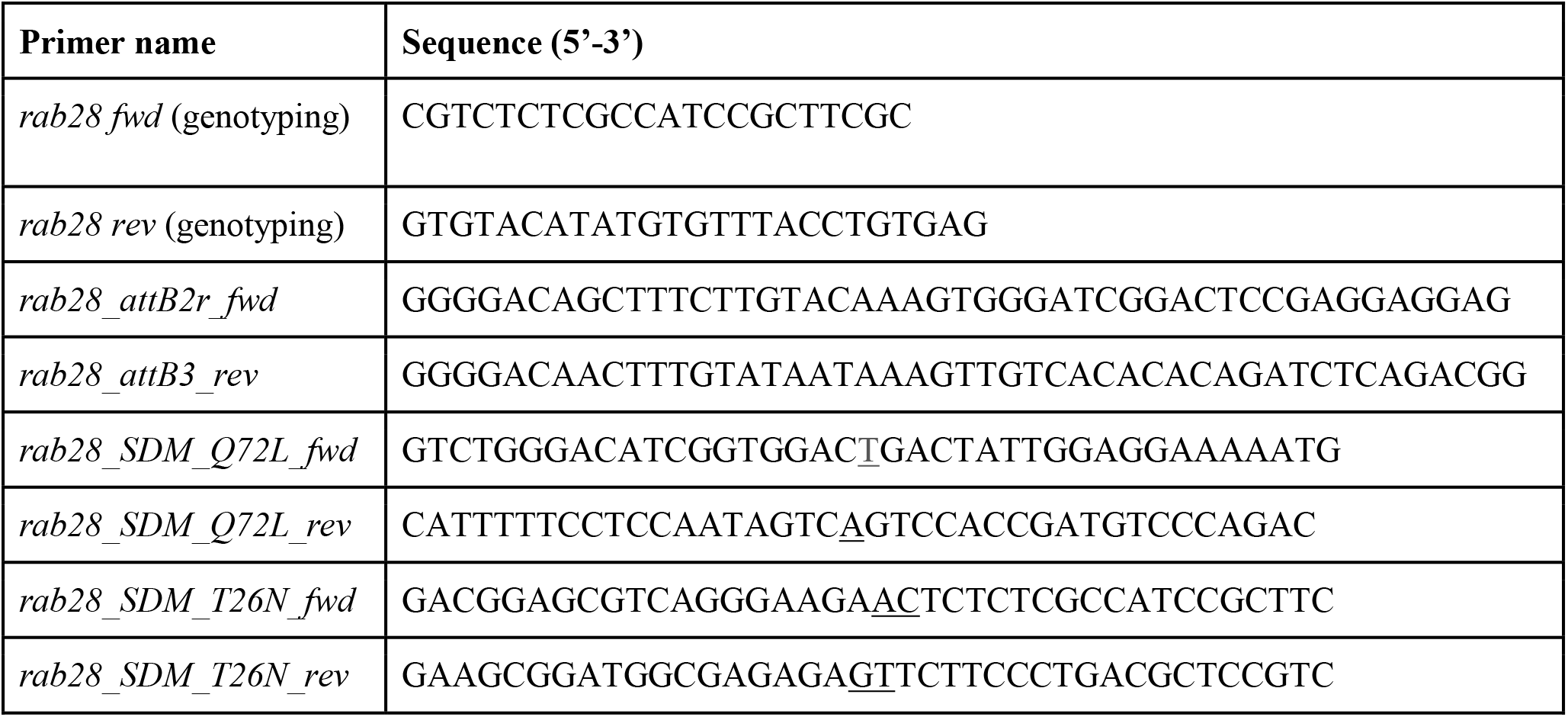
Sequences of primers used in this study. Underlined nucleotides indicate mutated positions.

### Behavioural assays

The optokinetic response (OKR) assay was performed by immobilising 5 dpf larvae in 9% methylcellulose in a 55 mm petri dish. The dish was placed inside a rotating drum with a black and white striped pattern with 99% contrast on the inside. The drum was rotated clockwise and anticlockwise for 30 sec each, at a speed of 18-21 rpm, during which time the number of eye movements (saccades) of the fish were manually recorded using a stereomicroscope. The visual motor response (VMR) assay was performed using the ZebraBox^®^ recording chamber (ViewPoint). 5 dpf larvae were placed in individual wells of a 96 well polystyrene plate in 600 μl of embryo medium, which was placed in the recording chamber. Locomotor activity of the larvae in response to changing light conditions was recorded using an infrared camera. Data analysis was performed as previously described (Deeti et al. 2014).

### Protein extraction and Immunoblotting

5 dpf zebrafish larvae were killed on ice and eyes dissected in a solution of 5 mM NaCl with protease inhibitor cocktail tablets (Roche). Eyes were either snap frozen in liquid nitrogen and stored at -80°C or immediately lysed. Protein concentration was estimated by Bradford assay to ensure equivalence between samples. Proteins were separated on a 0.75 mm 12% Bis/Tris acrylamide SDS-PAGE resolving gel and transferred to nitrocellulose membranes. Membranes were blocked in 5% skim milk-PBST for 1 hour at room temperature and subsequently probed with primary antibodies at 4°C overnight, followed by secondary antibodies. Primary antibodies used in this study were anti-GFP (1:500, Santa Cruz Biotechnology) and anti-PDE6D (1:500, Abcam). Secondary antibodies were HRP-conjugated anti-rabbit or HRP-conjugated anti-mouse (both 1:2000, Cell Signalling Technology).

### Immunoprecipitation

Immunoprecipitations were performed on 5 dpf larval eyes (100 per replicate) lysed in IP lysis buffer (50 mM Tris HCl, pH 7.5, 150 mM NaCl, 10 mM MgCl_2_, 1% NP-40, 1 mM DTT, 1 mM PMSF, 2 mM Na_3_VO_4_ and protease inhibitor cocktail (Roche), 1 tablet per 10 ml). Tissue was disrupted by aspiration through a needle and syringe, followed by a 20 min incubation on a tube rotator (Stuart) at 4°C. Lysates were centrifuged at 20,000 x g for 15 min, the supernatant was loaded onto GFP-Trap beads (Chromotek) and incubated on a rotor at 4°C for 2 hrs or overnight. Following this the beads were pelleted by centrifugation at 2500 g and washed three times with lysis buffer. For immunoblotting, proteins were eluted from the beads with SDS sample buffer followed by boiling at 95°C for 5 min. For mass spectrometry, a previously described protocol was followed (Turriziani et al. 2014). Briefly, proteins were trypsinised on the beads in 60 μl of Buffer I (2 M urea, 50 mM Tris-HCl pH 7.5, 5 μg/ml Trypsin [modified sequencing-grade trypsin; Promega]) for 30 min at 37°C in a thermomixer, shaking at 700 rpm. Samples were briefly centrifuged and supernatants transferred to clean Eppendorf tubes. The beads were then incubated in 50 μl Buffer II (2 M urea, 50 mM Tris-HCl pH 7.5, 1mM DTT) for 1 h at 37°C, shaking at 700 rpm in a thermomixer. Samples were again briefly centrifuged, and the supernatants from Buffers I and II pooled and left to continue trypsin digestion overnight at room temperature.

### Mass spectrometry

Samples from the overnight digest were alkylated by addition of 20 μl iodoacetamide (5 mg/ml) and incubation for 30 min in the dark. 1 μl 100% trifluoroacetic acid (TFA) was added to the samples to stop the reaction and samples were then loaded onto equilibrated C18 StageTips containing octadecyl C18 disks (Sigma) (Turriziani et al. 2014). Briefly, a small disc of Empore material 3M was inserted into a pipette tip, preparing a single tip for each sample. Tips were activated and equilibrated by washing through 50 μl of 50% acetonitrile (AcN) – 0.1% TFA solution followed by 50 μl of 1% TFA solution, using a syringe to pass liquid through the pipette tips. Once added to StageTips, samples were desalted by washing twice with 50 μl of 1% TFA solution. Peptides were then eluted into clean Eppendorf tubes using 2 x 25 μl 50% AcN – 0.1% TFA solution. The final eluates were concentrated in a CentriVap concentrator (Labconco, USA) and re-suspended in 12 μl 0.1% TFA solution, ready for analysis by mass spectrometry (Turriziani et al. 2014). Peptides were analysed on a quadrupole Orbitrap (Q-Exactive, Thermo Scientific) mass spectrometer equipped with a reversed-phase NanoLC UltiMate 3000 HPLC system (Thermo Scientific). To identify peptides and proteins, MS/MS spectra were matched to the UniProt Danio rerio database. LFQ intensities were subsequently analysed using Perseus (v1.6.1.3) (Tyanova et al. 2016). Protein identifications were filtered to eliminate the identifications from the reverse database and common contaminants. Data was log2 transformed and t-test comparison of fractions carried out. Gene ontology terms were identified and visualised by submitting identified gene lists to the PANTHER database (Thomas et al. 2003).

### Fluorescence microscopy

Zebrafish larvae were fixed in 4% PFA overnight at 4°C and subsequently washed with PBS, cryoprotected in a sucrose gradient ascending series and finally embedded in OCT (VWR). 10 μm thick frozen sections were cut on a Microm HM 505 E cryostat and mounted on Superfrost Plus slides (ThermoFisher). Sections were stained with the following primary antibodies: rat anti-GFP (1:500, Santa Cruz Biotechnology), rabbit anti-UV opsin (1:250, a gift from David Hyde (Vihtelic et al. 1999)) or rabbit anti-cone transducin α (1:50 a gift from Susan Brockerhoff (Brockerhoff et al. 2003)). Secondary antibodies were Alexa 488 or Alexa 567-conjugated (1:500, Thermo Fisher), respectively. Following antibody incubation, sections were stained with DAPI. Slides were then mounted with Mowiol^®^ (Merck) and cover slipped.

Immunostained zebrafish retinal sections were imaged on an inverted Zeiss LSM 510 Meta confocal laser scanning microscope. High-resolution images of eGFP-Rab28 localisation were taken with an Olympus FLUOVIEW FV3000 confocal microscope for 5 dpf retinas and with a Leica TCS SP8 X for 1 mpf retinas (resolution 120-200 nm). Images were deconvolved using Huygens Professional software (Scientific Volume Imaging B.V) All image analysis was performed using Fiji (Schindelin et al. 2012).

### Transmitted light microscopy

Zebrafish were euthanised with tricaine methanesulfonate and the eyes enucleated and fixed overnight at 4°C in 2% PFA and 2.5% glutaraldehyde in 0.1 M Sorenson phosphate buffer pH 7.3. Samples were post-fixed in 1% osmium tetroxide and dehydrated in a gradient ascending series of ethanol concentrations prior to Epon 812 resin embedding overnight. 1 μm sections were prepared using a Leica EM UC6 microtome and glass knife, mounted on glass slides and stained with toluidine blue. The appearance of the lens core was used as a landmark to ensure similarity of samples in imaging and measuring. Sections were imaged with a Nikon eclipse 80i upright microscope equipped with a Canon EOS 600D camera.

### Transmission Electron Microscopy

Zebrafish eyes were embedded for TEM using the same protocol for light microscopy. 90 nm sections were cut on a Leica EM UC6 microtome, mounted on copper grids and post-stained with 2% uranyl acetate and 3% lead citrate. Imaging was performed on an FEI Tecnai 120 electron microscope.

## Results

### Zebrafish *rab28* and CRISPR mutagenesis

We initiated CRISPR knockouts by characterising the zebrafish orthologue of *rab28*. While the locus and genomic sequence were unknown, a sequenced mRNA transcript was reported (RefSeq: NM_199752.1). Human RAB28 has three splice isoforms (Roosing et al. 2013), whereas there is only one known zebrafish isoform. It most closely matches the human RAB28S isoform in sequence (**Figure 1A**). Due to poor annotation of the zebrafish *rab28* locus, it was necessary to obtain genomic DNA sequence to accurately design CRISPR sgRNAs to target the gene. To design primers for sequencing it was necessary to estimate the exon-intron structure of zebrafish *rab28*. Primers were designed to each potential exon of *rab28* to amplify the intervening introns. A product was successfully amplified in three reactions, the others failing either because the prediction of exon placement was incorrect or the introns were too large to amplify by PCR (**Figure 1B**). The amplified products were subcloned and DNA sequencing confirmed the positions of exons 2 and 3 as predicted (**Figure 1C and D**). This sequence information facilitated design of sgRNAs and genotyping strategies for the *rab28* KO lines. Cas9-rab28exon2 sgRNA ribonucleoprotein particles (RNPs) were injected into one-cell stage embryos and raised to adulthood. These adult P_0_ fish were genotyped for the presence of mutant *rab28* alleles by PCR and outcrossed to enable germline transmission. Two *rab28* mutant lines were generated, *rab28^ucd7^* and *rab28^ucd8^*, a 40 bp deletion and a 14 bp insertion or a 65 bp deletion and an 8 bp insertion, respectively. Both alleles disrupt part of exon 2 coding sequence and the intron 2 donor site. Retention of intron 2 is predicted to lead to truncation of Rab28 due to the presence of in-frame stop codons and therefore the loss of several functional motifs from the translated protein (**Figure 1D**). RT-PCR of RNA from homozygous knockout larvae shows an absence of the correctly spliced transcript (**Figure 1E**), suggesting that the transcript is probably degraded by nonsense-mediated decay. Zebrafish homozygous for each of these alleles were subsequently used in phenotypic analyses.

**Figure 1.**
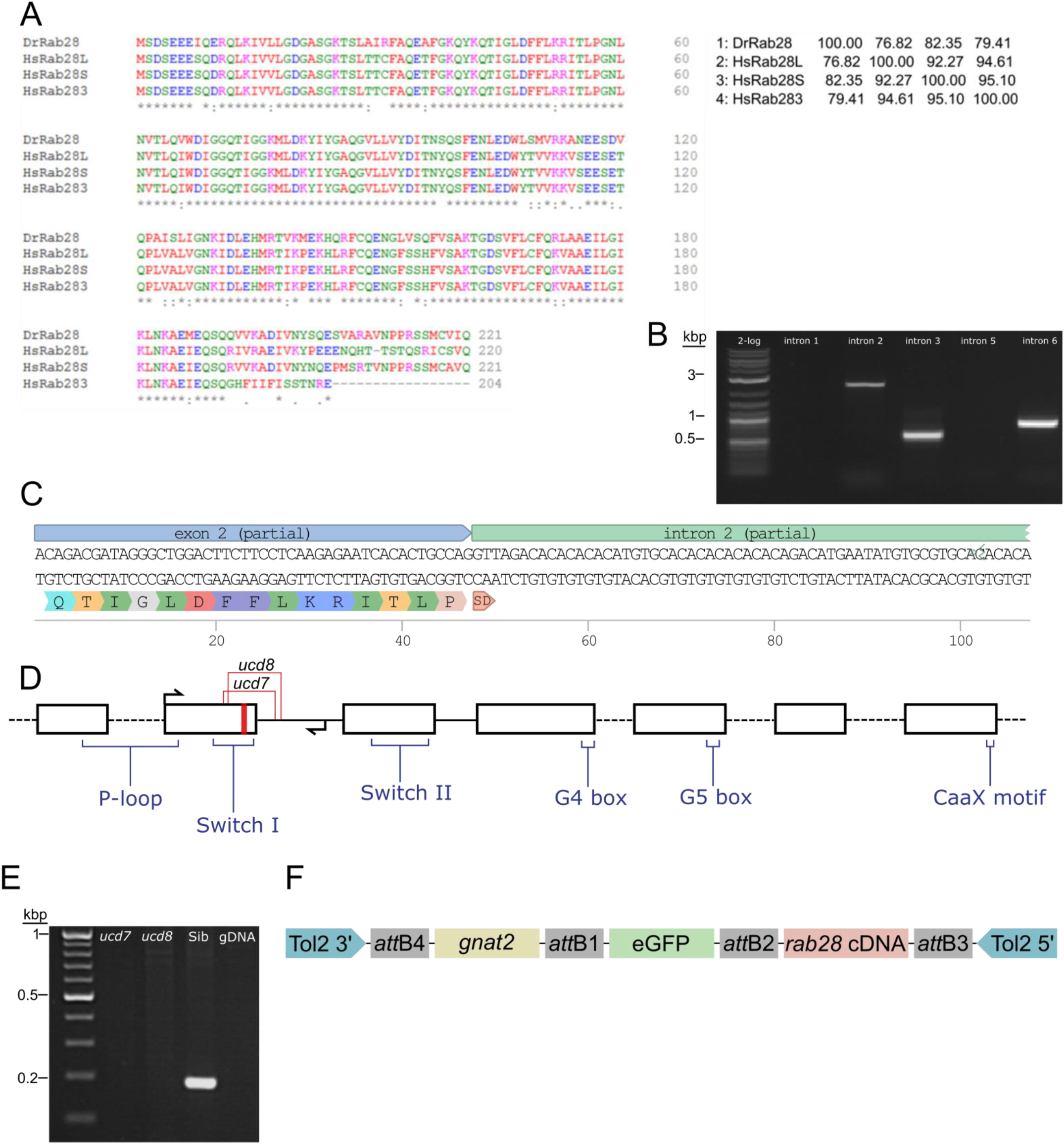
Sequencing of the zebrafish *rab28* gene, CRISPR mutagenesis and Tol2 transgenesis. **(A)** Multiple sequence alignment of zebrafish Rab28 protein sequence, predicted from mRNA, with that of the three known human RAB28 protein isoforms. The percentage protein identity is shown in a matrix table. **(B)** Representative gel showing PCR amplification of predicted *rab28* introns 2, 3 and 6. **(C)** DNA sequencing reveals the exonintron boundary at exon 2 of zebrafish *rab28* and the predicted translated sequence encoded by exon 2. SD: splice-donor site. **(D)** Schematic of predicted *rab28* gene structure. Black boxes represent exons and connecting lines represent introns. Red stripe indicates location of sgRNA target site used to generate CRISPR mutants, arrows indicate genotyping primer positions, coding positions for critical Rab GTPase motifs are highlighted and the location of *ucd7* and *ucd8* indels indicated. **(E)** Example RT-PCR gel showing the absence of correctly spliced *rab28* cDNA between exons 2 and 4 in homozygous *ucd7* and *ucd8* larvae, which is present in WT siblings. **(F)** Schematic of the construct used to generate eGFP-Rab28 transgenic zebrafish. Expression is driven by the *gnat2* (cone transducin alpha) promoter. *att:* Gateway att sites, Tol2 3’ and 5’: transposon inverted repeats.

In order to assess localisation, function and protein-protein interactions of Rab28, we also generated a transgenic fish line expressing an eGFP-Rab28 construct in cone photoreceptors (**Figure 1F**). We also generated two further transgenic lines, one harboring the (predicted GTP-preferring) Q72L mutation and another the (predicted GDP-preferring) T26N mutation. It should be noted that the TN mutation commonly used to ‘GDP-lock’ Rabs lowers the affinity for guanine nucleotides generally (Lee et al. 2009), so our T26N mutant may also mimic the nucleotide empty state.

### eGFP-Rab28 localisation to cone outer segments is partially dependent on GTP/GDP binding

GTPase switching between the GTP or GDP-bound conformations is often accompanied by a change in protein localisation to another cellular compartment. We previously reported that GTP and GDP-binding variants of *C. elegans* RAB-28 dramatically alter localisation in ciliated sensory neurons (Jensen et al. 2016). While RAB28 localises to murine rod and cone OS (Ying et al. 2018), it is unknown if this is influenced by nucleotide binding. This question was investigated with three eGFP-Rab28 variants expressed in zebrafish cones. Confocal imaging of cryosections from 5 dpf zebrafish revealed eGFP-Rab28 localised almost exclusively to cone OS (**Figure 2A-C**). However, in the T26N mutant, an average 30% reduction in OS enrichment was observed compared to the WT variant (**Figure 2D;** *p*<0.0001), suggesting less efficient targeting of GDP-bound or nucleotide empty eGFP-Rab28 to OS. Intriguingly, the eGFP-Rab28^T26N^ mutant reporter was observed to occasionally localise to discrete, horizontal bands in the OS of some cones (Figure 2E-F). The OS of zebrafish photoreceptors are fully mature by 24 dpf (Branchek & Bremiller 1984). To investigate whether further photoreceptor development is accompanied by changes in Rab28 localisation, retinal sections were imaged from 1 mpf zebrafish. Again, the WT and mutant versions of eGFP-Rab28 were strongly enriched in the OS of all cones (Figure 3A-C). Strikingly, at this time point, discrete banding patterns were observed in all three transgenic lines, in a larger number of photoreceptors and was far more extensive than in larvae, occurring at regular intervals from base to tip of cone OS (Figure 3-F). In the WT and Q72L mutant reporters, banding appeared largely restricted to the short single (SS) cone population, located in the bottommost row of photoreceptors (Figure 3A,B,D and E). The T26N reporter, by contrast, showed discrete banding in other cone populations (Figure 3C and F). Our data show that Rab28 is efficiently targeted to cone OS in a manner only partially dependent on its nucleotide-bound state, where it is organised into discrete segments of the OS, a behaviour that appears to be more prominent when in the GDP-bound/nucleotide free state.

**Figure 2.**
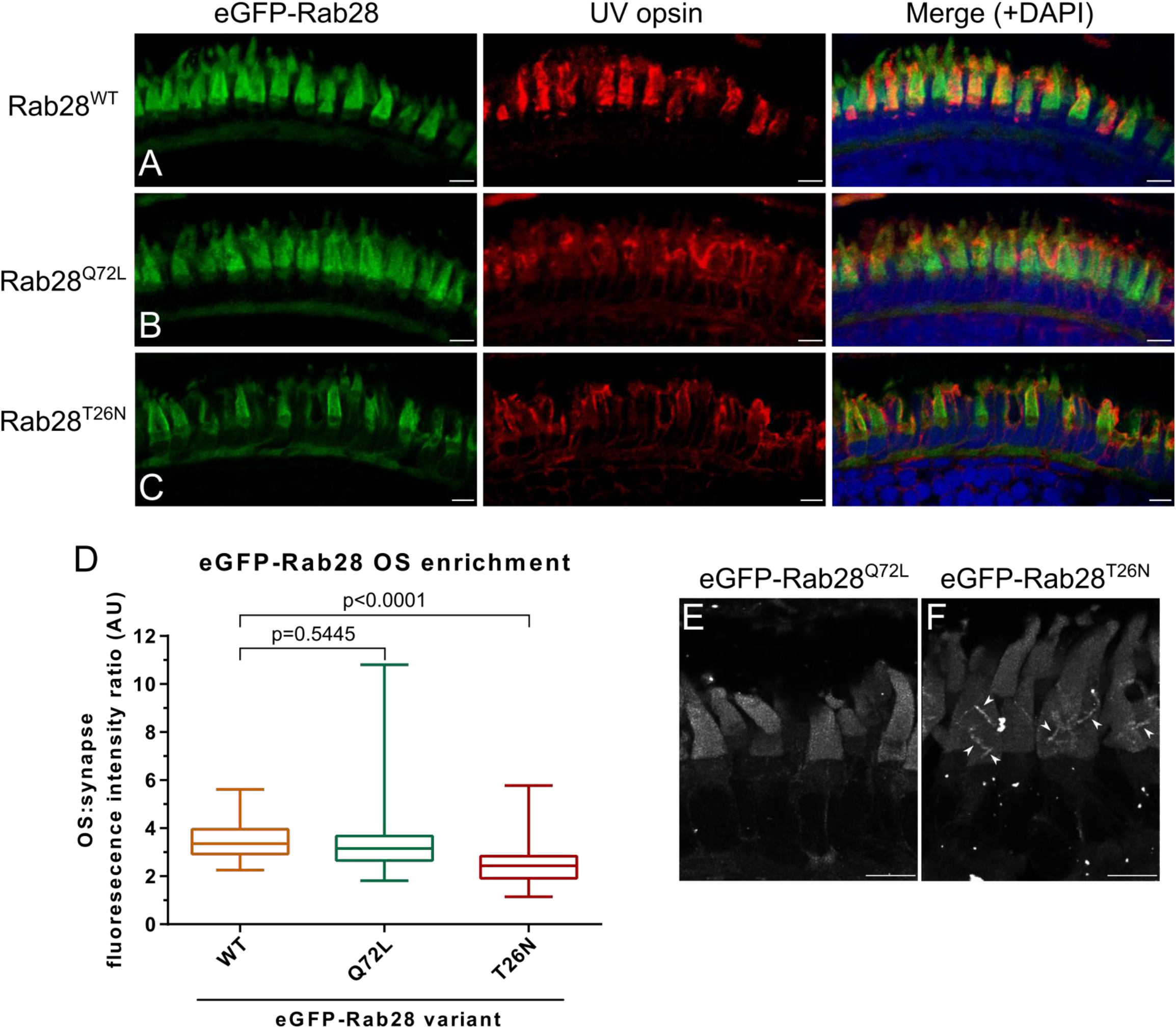
eGFP-Rab28 localisation to zebrafish cone outer segments is partially dependent on GTP/GDP-binding. **(A-C)** Representative confocal z-projections of eGFP-Rab28 localisation in 5 dpf zebrafish cone photoreceptors. The WT, putative GTP-preferring (Q72L) and GDP-preferring (T26N) variants of Rab28 all localise strongly to the outer segments of zebrafish cones, co-localising with UV opsin labelling. Scale bars; 5 μm. For WT, Q72L and T26N eGFP-Rab28 reporters a total of 14, 24 and 25 larvae were imaged, respectively. **(D)** Box and whisker plots of the ratio of eGFP-Rab28 intensity in the OS vs synaptic region of larval cones. Box extremities represent 1st and 3rd quartiles; whiskers are maximum and minimum values. Data are from 60 cones per transgenic line. One-way ANOVA p-value < 0.0001. **(E-F)** Deconvolved, high resolution confocal z-projections of eGFP-Rab28 Q72L and T26N mutant localisation in cones of 5 dpf larvae. A discrete localisation pattern of the T26N mutant in COS is clearly observed (white arrowheads). Scale bars 4 μm.

**Figure 3.**
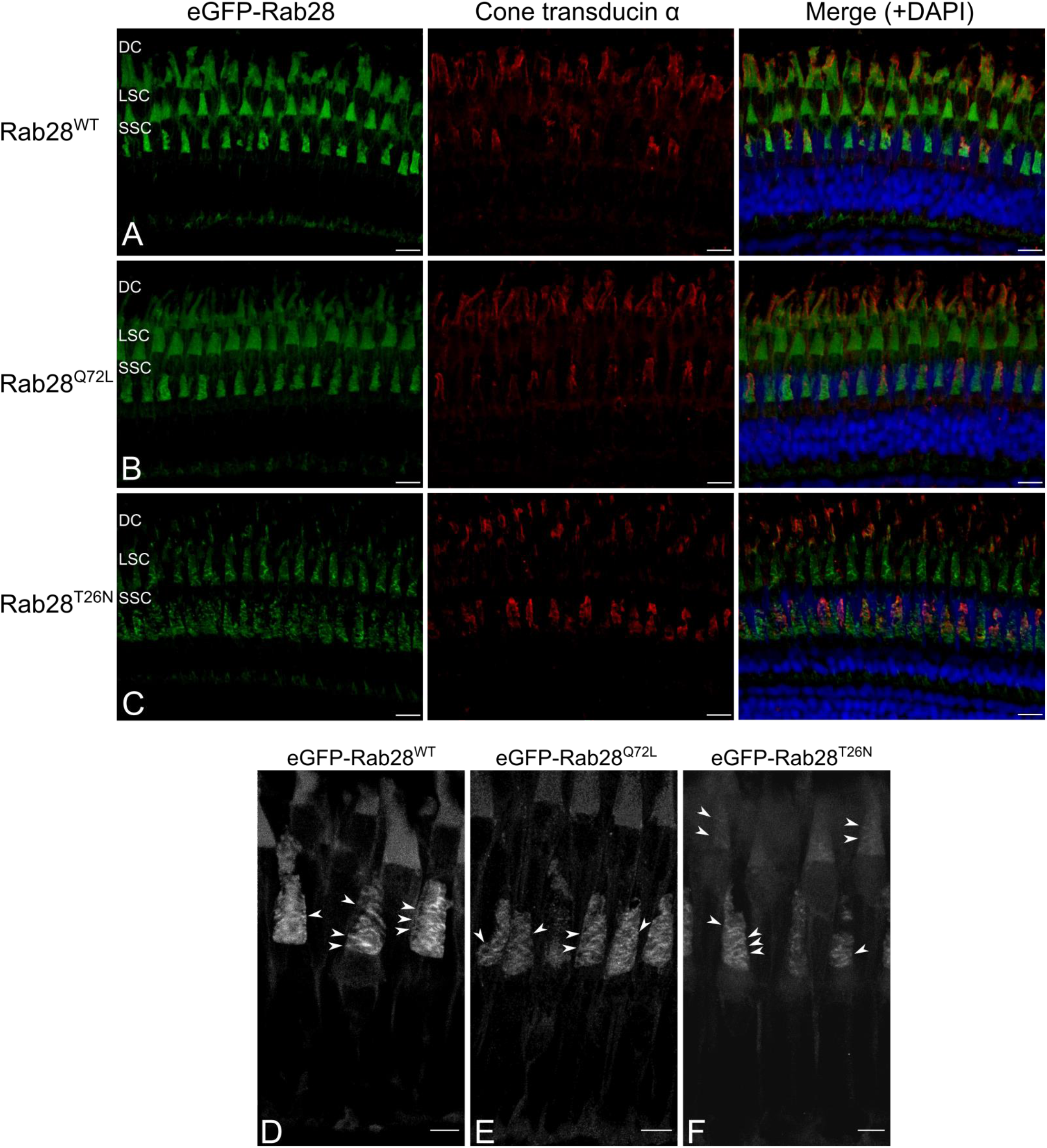
eGFP-Rab28 localisation in 1 mpf cone photoreceptors. **(A-C)** Representative confocal z-projections of eGFP-Rab28 localisation in 1 mpf zebrafish cones. Cone OS are stained with anti-cone transducin alpha antibody. Each tier of photoreceptors is comprised of different classes of cone. DC: double cones; LSC: long single cones; SSC: short single cones. Scale bars 10 μm. For WT, Q72L and T26N eGFP-Rab28 reporters a total of 13, 11 and 11 retinas were imaged, respectively. **(D-F)** Deconvolved, high resolution confocal z-projections of eGFP-Rab28 WT, Q72L and T26N mutant localisation in 1 mpf cones. Arrowheads point to discrete bands present throughout the OS. Scale bars 5 μm.

### *rab28* mutant zebrafish have normal visual function at 5 dpf

Compared to sibling controls, *rab28*^-/-^ larvae display normal development and gross morphology at 5 dpf (**Figure 4A**). To assess visual function in *rab28*^-/-^ zebrafish, we utilised two behavioural assays: the optokinetic response (OKR) and visual-motor response (VMR). Homozygous mutant larvae and control (+/+ and +/-) siblings for both CRISPR alleles were assessed at 5 dpf. The OKR of *rab28*^-/-^ larvae was not different to control siblings at 5 dpf (**Figure 4B**). To investigate the possibility of a reduction in visual function at a later age, we performed OKR assays on 21 dpf *rab28* mutants and controls (**Supplementary figure 1A**). As at 5 dpf, the OKR of *rab28* knockouts was not significantly different from siblings at 21 dpf (*p*=0.5224). In the VMR assay, the OFF peak activity between siblings and mutants is identical, whereas *rab28* mutants display a 51% increased average ON peak activity (**Figure 4C**; *p*=0.0017). The overall activity traces and the OFF and ON peak traces (100 s before and 400 s after a light change) highlight *rab28* mutants with slightly elevated activity in the dark, but reduced activity in light conditions, compared to sibling controls (**Figure 4D**). These data show that, in zebrafish at 5dpf, *rab28* knockout results in subtle adverse effects on visual behaviour.

**Figure 4.**
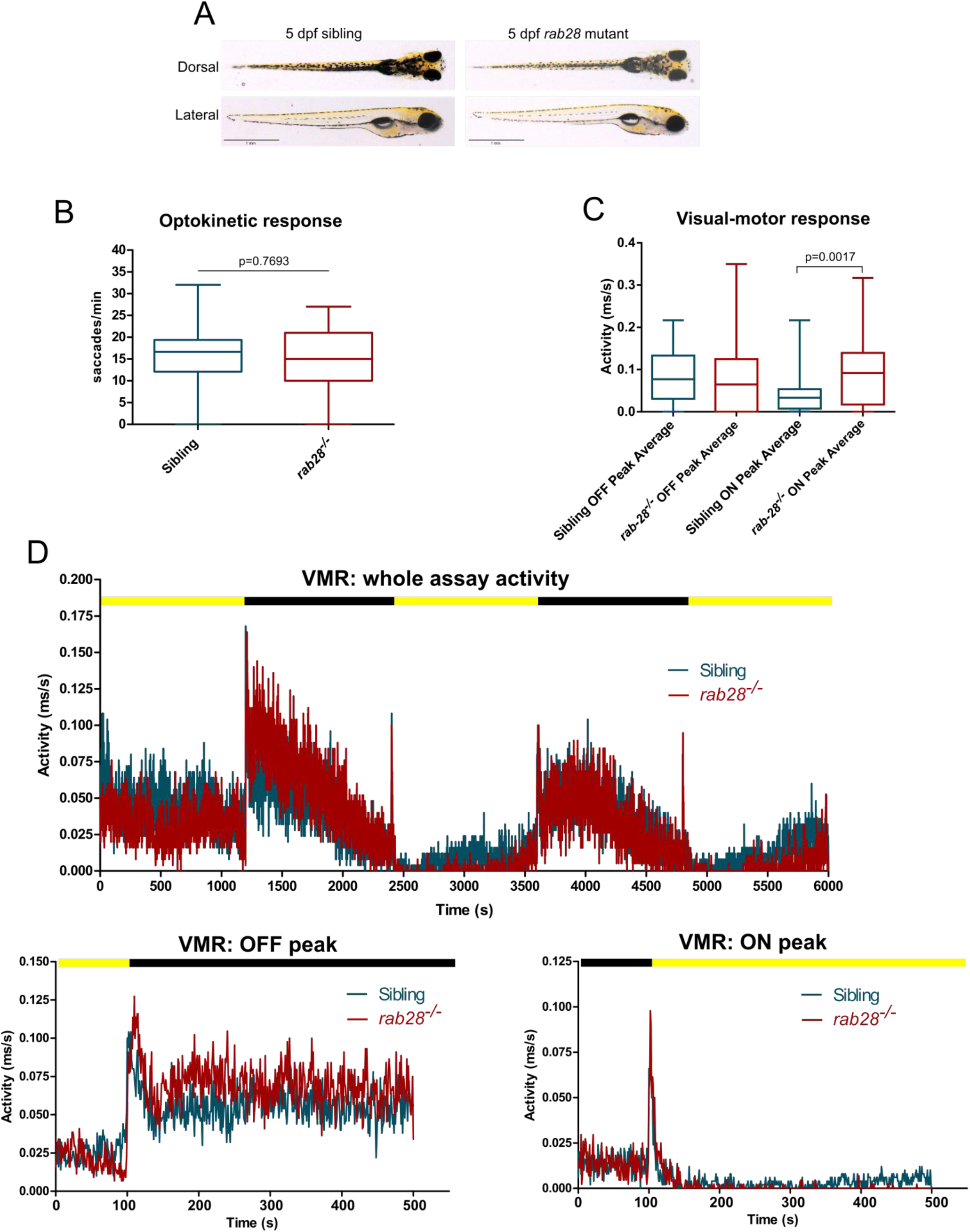
*rab28* knockout zebrafish have subtle defects in visual behaviour at 5 dpf. **(A)** Representative images of *rab28* knockout and sibling zebrafish larvae at 5 dpf. Gross morphology is indistinguishable between knockouts and siblings. (B) Box and whisker plot of the optokinetic response (OKR) of *rab28* knockout larvae versus sibling larvae at 5 dpf. OKRs are not significantly different. Box extremities represent 1st and 3rd quartiles; whiskers are maximum and minimum values. p-value derived from unpaired t-test. OKR data are from 32 mutants and 98 siblings, across three experimental replicates. **(C)** Scatter plots of 5 dpf larval activity during the visual-motor response (VMR) assay. OFF peak activity is identical between *rab28* knockouts and siblings, albeit *rab28* knockouts have an average 51% higher ON peak activity. Error bars show SEM, p-value derived from unpaired t-test. **(D)** Activity traces showing 5 dpf larval activity over the course of an entire VMR assay (100 min), as well as separate graphs showing activity 100 s before and 400 s after OFF and ON peaks, respectively. Black and yellow bars indicate dark and light conditions, respectively. VMR data are from 32 mutants and 49 siblings, across three experimental replicates.

### eGFP-Rab28 transgenic zebrafish have reduced visual function at 5 dpf

We previously demonstrated overexpression of either GTP or GDP-preferring RAB28 induces functional and ultrastructural defects in the cilia and sensory organs of the nematode *C. elegans*. To assess an evolutionarily conserved function of this nucleotide binding domain in the vertebrate retina, we assessed visual function in transgenic eGFP-Rab28 zebrafish larvae (**Figure 5A-E; Supplementary figure 1A-E**). In the OKR, 5 dpf transgenic larvae expressing eGFP-Rab28^WT^ displayed normal saccadic eye movements equivalent to non-transgenic fish (18-25/min) (**Figure 5A**). eGFP-Rab28^Q72L^ larvae, however, had a far greater range of responses and an average 30% reduction in OKR, while the eGFP-Rab28^T26N^ expressing larvae had similar responses to WT (**Figure 5A**).

**Figure 5.**
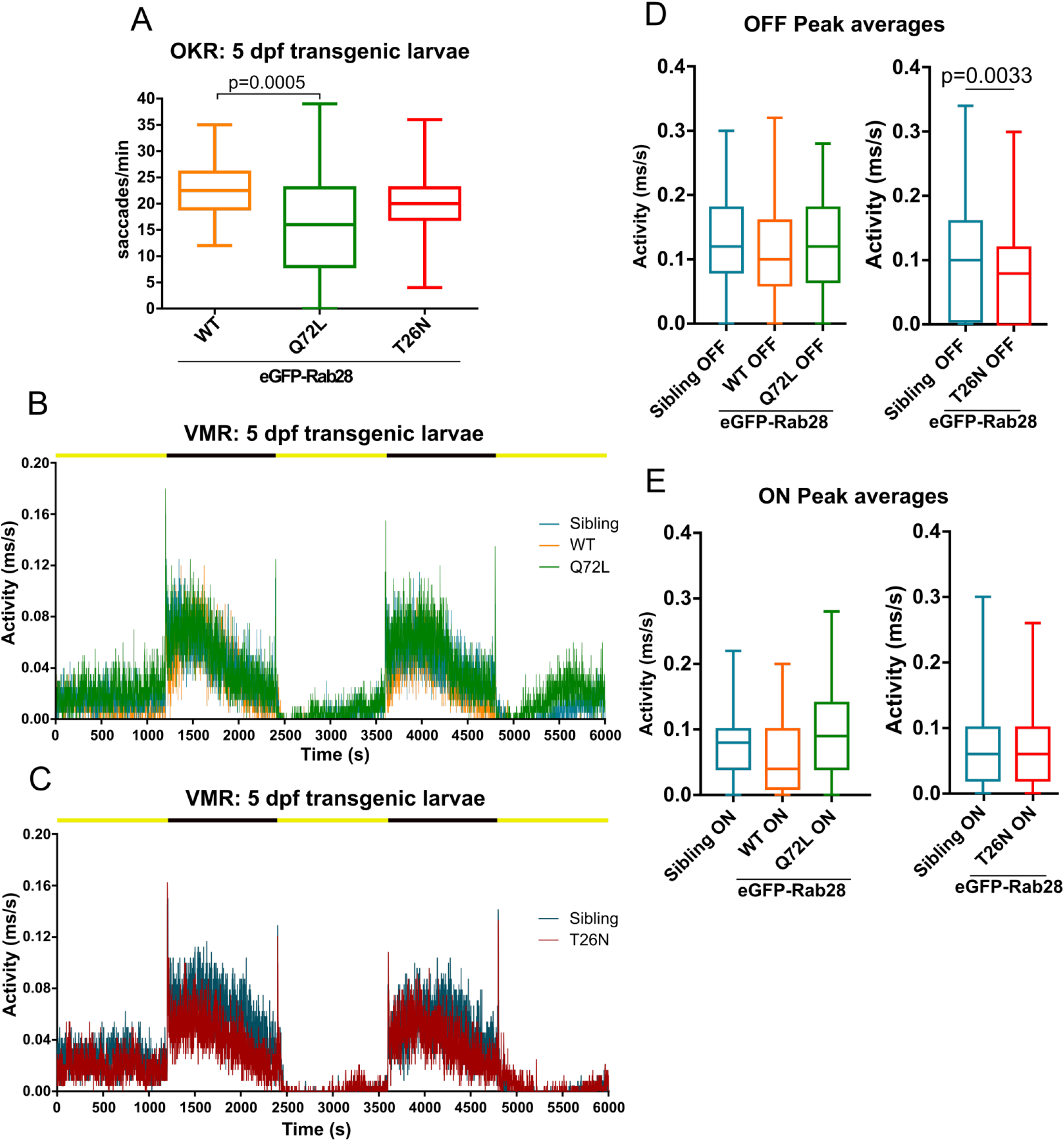
Rab28 transgenic zebrafish have mild to moderate visual defects at 5 dpf. **(A)** Box and whisker plot of optokinetic response (OKR) assay of 5 dpf larvae overexpressing GFP-Rab28 WT, Q72L (GTP-preferring) or T26N (GDP-preferring). Larvae expressing the Q72L variant display greater variability and an overall reduction in OKR scores. Box extremities represent 1st and 3rd quartiles; whiskers are maximum and minimum values. Data are from three independent replicates; at least 30 larvae analysed per strain over three experimental replicates; p-value derived from one way ANOVA. (**B** and **C**) Representative activity traces of 5 dpf transgenic and sibling larval activity over the course of the entire VMR assay. Black and yellow bars indicate dark and light conditions, respectively. (**D** and **E**) Box and whisker plots of the OFF and ON peak activity of 5 dpf transgenic and sibling larvae. Data are from three independent replicates and are the average of 5 seconds of activity following light changes. At least 64 larvae were analysed per strain. p-value derived from t-test **(D)**.

By contrast, the VMR assay of eGFP-Rab28^WT^ and eGFP-Rab28^Q72L^ 5 dpf larvae showed similar light and dark activity compared to non-transgenic sibling controls (**Figure 5B,D and E; Supplementary figure 1B and C**), while the dark, but not light, activity of eGFP-Rab28^T26N^ larvae was reduced (**Figure 5C, D and E; Supplementary figure 1D and E**). T26N transgenic larvae also had significantly reduced OFF peak activity compared to siblings (**Figure 5D**). Overall, these data show that transgenic larvae overexpressing eGFP-Rab28 GTP and GDP-preferring mutants display mild to moderate defects in visual function at 5 dpf.

### *rab28* mutants have normal retinal histology and ultrastructure up to 12 mpf

The absence of visual behaviour deficits in larval and juvenile *rab28*^-/-^ fish led us to investigate the possibility of a slow-onset, progressive retinal degeneration, as observed in other zebrafish models, such as *eys* and *rpgrip1* (Yu et al. 2016; Raghupathy et al. 2017). Thus, homozygous *rab28*^-/^ larvae were raised to adulthood and retinal histology assessed. At 3 mpf, the retina of a *rab28*^-/-^ had equivalent retinal lamination to a sibling control and the photoreceptor layer contained all five photoreceptor cell types in their normal distribution and abundance (**Figure 6A**). We then assessed retinal histology in a 12 mpf sibling and *rab28* mutant and again found mutant retinas to be healthy, with no evidence of degeneration. To investigate potential ultrastructural defects, TEM was performed on 3 mpf retinas. Photoreceptor ultrastructure was similar between sibling and mutant at 3 mpf. Both genotypes show normal basal body positioning, while cone lamellae in *rab28*^-/-^ fish display normal organisation and alignment (**Figure 6B**). Therefore, loss of *rab28* is not associated with pronounced retinal structure degeneration in zebrafish, up to 12 mpf.

**Figure 6.**
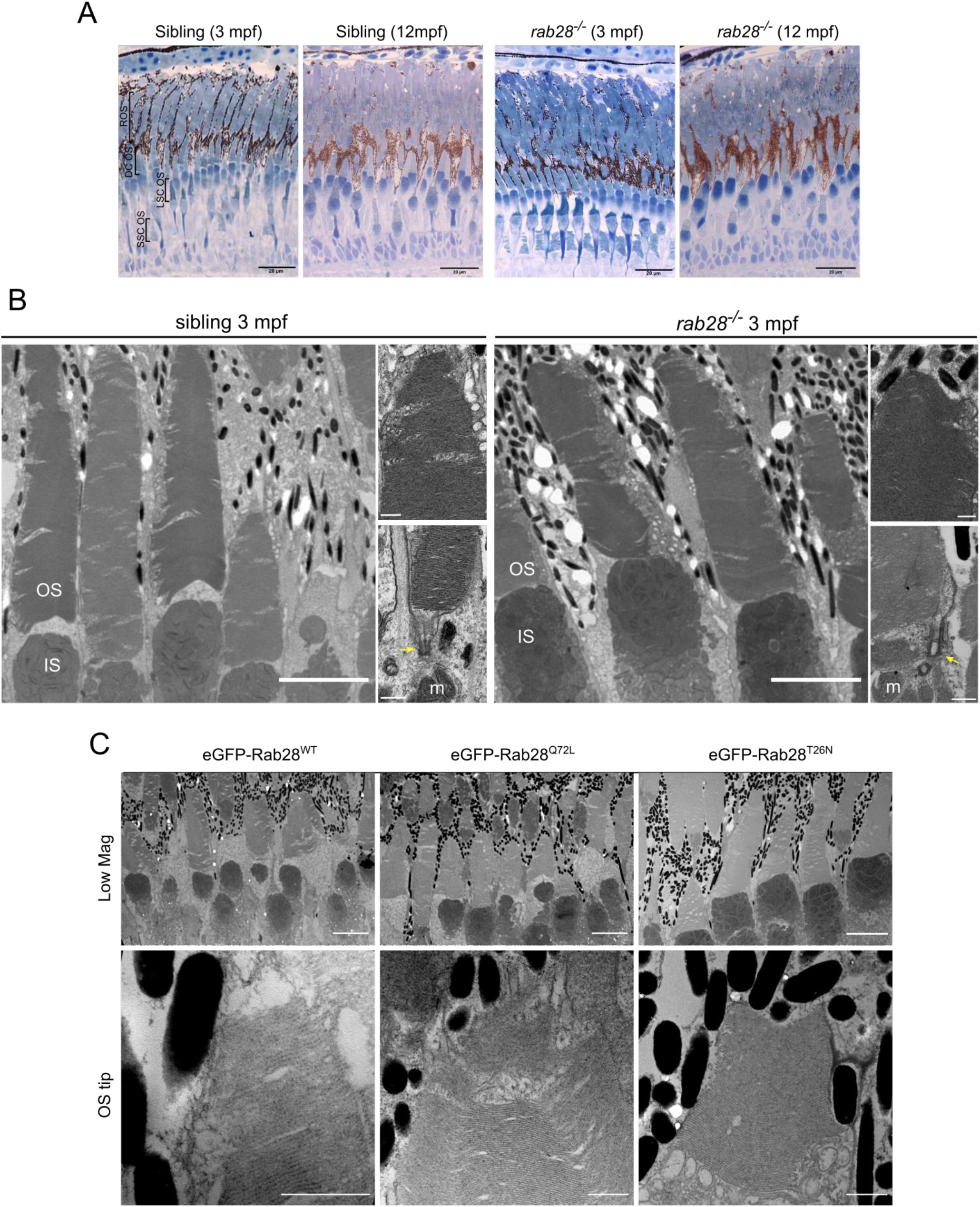
*rab28* knockout zebrafish have normal retinal histology and ultrastructure at 3 mpf. **(A)** Representative images of retinal histology in 3 mpf zebrafish, showing views of the photoreceptor layer in the central and peripheral retina in *rab28*^ucd8^ mutants and siblings. The overall structure and composition of the photoreceptor layer is grossly normal in both. Scale bars 20 μm **(B)** Representative transmission electron micrographs of 3 mpf zebrafish *rab28*^ucd7^ knockout and sibling retinas. Low magnification images show several cone photoreceptors, while high magnification images show examples of OS base and tips. Yellow arrows indicate ciliary basal body. OS: outer segment; IS: inner segment; m: mitochondria. Low magnification image scale bars 5 μm, high magnification scale bars 500 nm. **(C)** Representative TEM of 7 mpf eGFP-Rab28 transgenic zebrafish. Low magnification images show rows of several cone photoreceptors, while high magnification images show examples of OS tips. Low magnification scale bars 5 μm, high magnification scale bars 500 nm.

### eGFP-Rab28 transgenic zebrafish have normal photoreceptor ultrastructure at 7 mpf

Given the slight visual behaviour defects exhibited by eGFP-Rab28 mutant expressing larvae, we assessed photoreceptor ultrastructure in 7 mpf adults expressing the three different Rab28 reporters. We found that eGFP-Rab28 overexpressing cones had no obvious ultrastructural defects and normal outer segment morphology (**Figure 6C**). Thus, overexpression of eGFP-Rab28 or its GTP/GDP-preferring mutants does not adversely affect cone ultrastructure up to 7 mpf.

### *rab28* mutant zebrafish have reduced shedding of cone OS discs

Defective cone outer segment shedding was investigated in *rab28*^-/-^ zebrafish, as this phenotype was recently reported for *rab28*^-/-^ mice (Ying et al. 2018). Unlike mice, two shedding peaks are reported for zebrafish photoreceptors: one in the morning and one in the evening (Lewis et al. 2018). Both rods and cones shed at these time points. In *rab28*^-/-^ zebrafish, immunofluorescence staining was applied to identify cone OS protein staining (cone opsins and cone transducin alpha) located distal to the tips of the cone outer segments as a surrogate measure of RPE phagosomes containing shed outer segments. At 1-2 mpf, the number of cone phagosomes are reduced in *rab28* mutant zebrafish by ~43% compared to siblings (**Figure 7A-C**). For transgenic zebrafish, we used the eGFP-Rab28 itself to identify phagosomes. Surprisingly, all three transgenic models displayed normal levels of shedding (**Figure 7D-G**). Although there was a slight reduction in eGFP-Rab28^WT^ retinas, this was not statistically significant. Our data demonstrate a conserved role for Rab28 in OS shedding in zebrafish cones.

**Figure 7.**
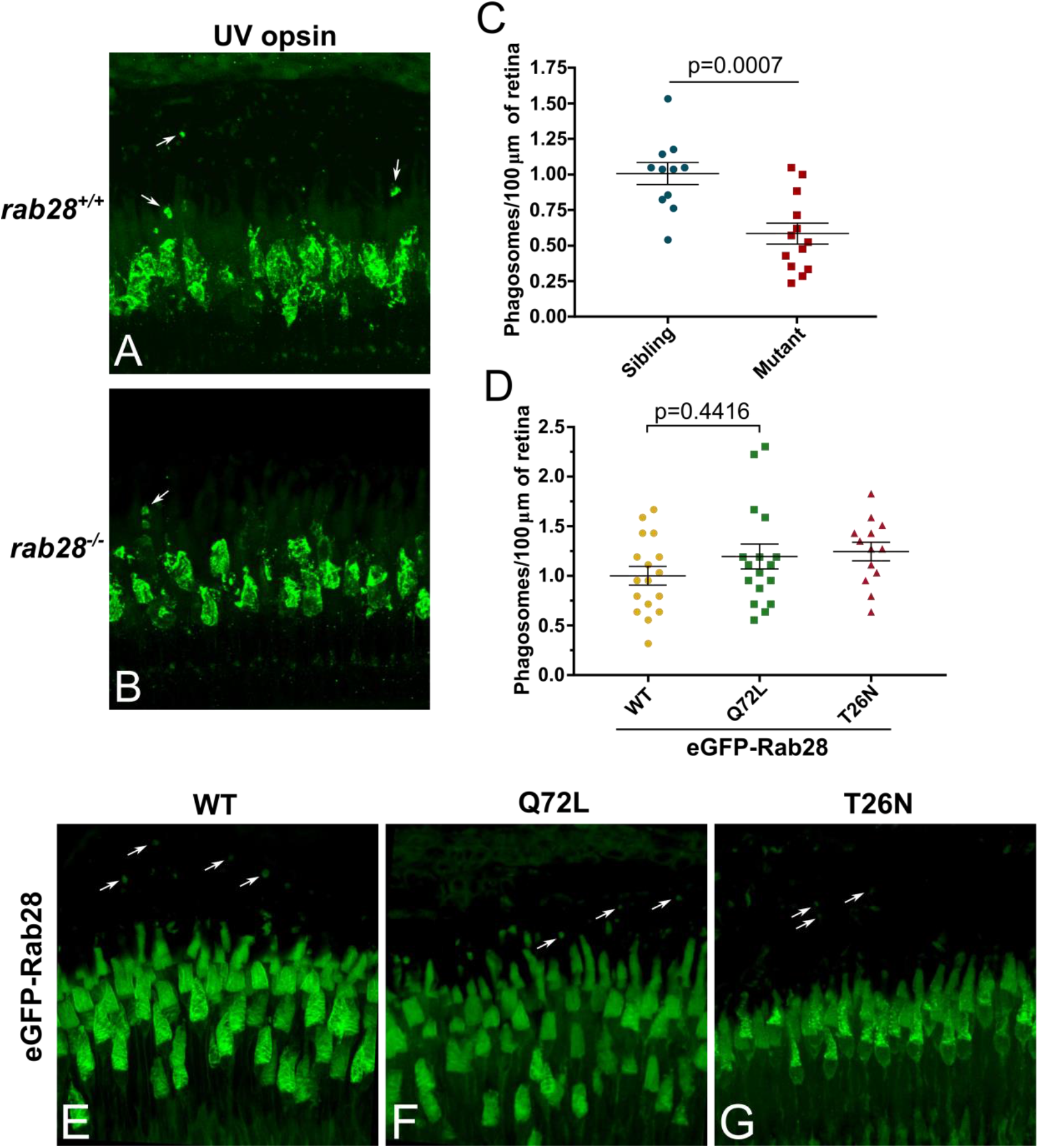
*rab28* knockout zebrafish have reduced outer segment shedding. **(A,B)** Confocal z-projections of *rab28* mutant and sibling control retinas at 1 mpf, stained for UV opsin to label phagosomes (white arrows). **(C)** Scatter plots of normalised phagosome density in *rab28* mutant and sibling retinas. p-value derived from t-test. Error bars show SEM. Data are from 13 and 11 retinal z-projections for mutant and sibling, respectively. **(D)** Scatter plots showing normalised phagosome density in the retinas eGFP-Rab28 transgenic zebrafish. p-value derived from one-way ANOVA. Error bars show SEM. **(E-G)** Confocal z-projections of 1 mpf retinas of zebrafish expressing eGFP-Rab28 (WT, Q72L or T26N variants). Phagosomes, labelled with eGFP, indicated by white arrows. Data are from 17, 17 and 13 retinal z-projections for WT, Q72L and T26N variants, respectively.

### Rab28 biochemically interacts with phototransduction proteins

In order to identify effectors and/or regulators of Rab28, immunoprecipitation (IP) of eGFP-Rab28 in 5 dpf zebrafish whole-eye lysates was performed (**Figure 8A**), followed by mass spectrometry. This was performed with eGFP-Rab28 WT, Q72L and T26N mutant lines, to identify interactants specific to the GTP and GDP-bound states. Using this approach, we identified 323 unique proteins across all three groups, of which 52 were deemed significantly enriched (t-test p<0.05) (**Figure 8B; Table 2; Supplementary Table 1**). The identified proteins can be divided into two groups based on fold change. The first group of 19 proteins have a log_2_ fold change >20 for at least one of the Rab28 variants, while the second group of 33 proteins have a log_2_ fold change <5 (**Table 2; Supplementary Table 1**), and cluster accordingly (**Supplementary figure 2**). To functionally categorise the Rab28 interactome, enriched gene ontology terms were identified using PANTHER-DB (**Figure 8C**). For the most enriched proteins across all three Rab28 variants, overrepresented processes and functions include signal transduction, cellular transport, metabolic processes and stimulus response (**Figure 8C**).

**Figure 8.**
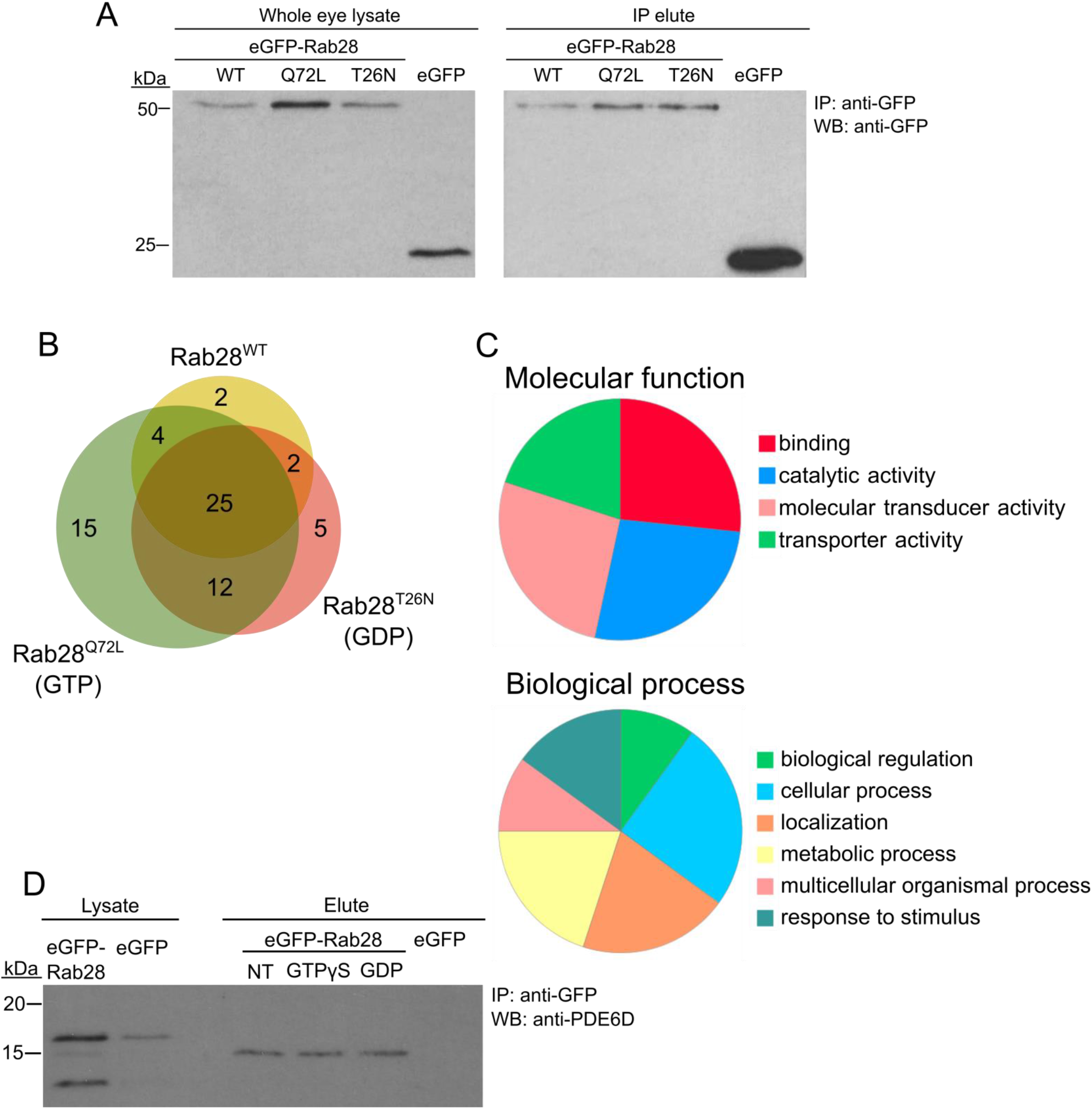
Rab28 interact with multiple phototransduction proteins in the zebrafish eye. **(A)** Western blotting with an anti-GFP antibody, showing eGFP-Rab28 in whole eye lysate and elute after immunoprecipitation with anti-GFP beads. An untagged eGFP only control is also shown. **(B)** Pie charts showing gene ontology terms for proteins identified by mass-spectrometry following co-immunoprecipitation, which demonstrate a statistically significant interaction with any of the three variants. **(C)** Venn diagram showing the number and overlap of significantly enriched (vs GFP-only control) interacting proteins identified for each variant. **(D)** Western blot of whole adult zebrafish eye lysate with an anti-PDE6D antibody, before and after IP of eGFP-Rab28. IPs either received no treatment, GTPγS or GDP. eGFP only control is also shown.

**Table 2:**
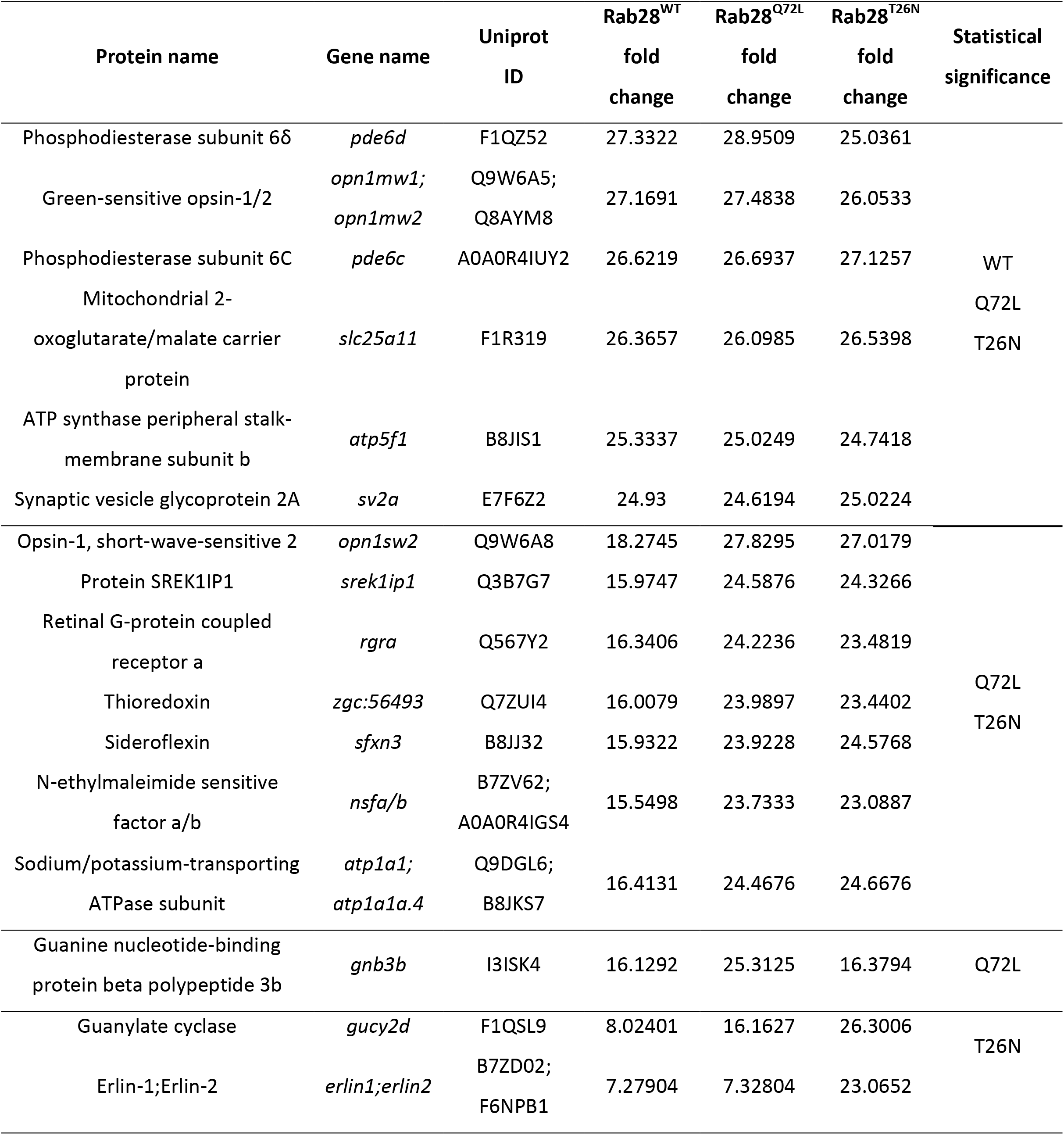
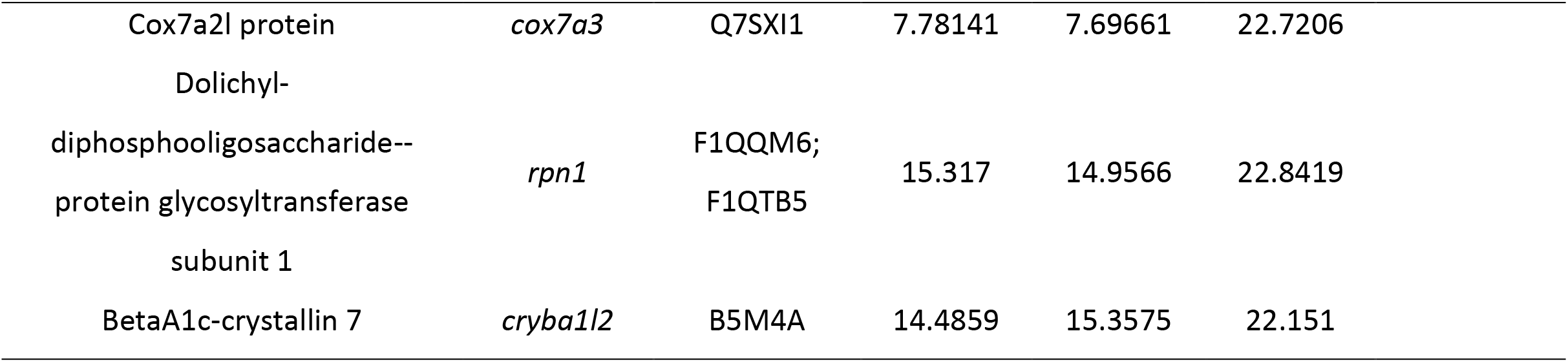
Selected interactant of RAB28 in the zebrafish eye. Shown are those proteins with a log_2_ fold change relative to the eGFP only control ≥ 20 for at least one of the eGFP-Rab28 variants.

There is significant overlap between the three Rab28 variants (**Figure 8B**), although a few proteins are enriched for one specific variant. Overall, the Rab28 interactome is highly enriched for components of the phototransduction cascade, including Green and Blue opsins, as well as Rhodopsin, Pde6c, Gucy2d and Cone transducin alpha (**Table 2; Supplementary Table 1**). Additionally, membrane transport proteins such as Nsfa/b (regulators of SNARE-mediated vesicle fusion), Sv2a and Erlin1/2 were significantly enriched. Unexpectedly, the interactome is also enriched for mitochondrial transport proteins (**Table 2; Supplementary Table 1**). The identification of some non-cone proteins (e.g. Rhodopsin, Rgra) is unsurprising, as the eGFP-Rab28 bait is exposed to potential interactants from other cell types in the whole-eye lysate.

Proteins specifically enriched for particular variants include Gnb3b, which is significantly enriched for the Q72L mutant alone, while Gucy2d and Erlin1/2 are significantly enriched for the T26N mutant only (**Table 2; Supplementary figure 2**).

One protein strongly detected across all three groups was the GDI-like solubilisation factor Pde6d, which is known to transport lipidated proteins to cilia and known to interact with Rab28 (Humbert et al. 2012; Ying et al. 2018). In our dataset, an equivalent fold change in Pde6d was detected across all three Rab28 groups (**Table 2**), although it was slightly higher for the Rab28^WT^ and Rab28^Q72L^ vs the Rab28^T26N^ (log2 FC = 27.33, 28.95 and 25.04, respectively), suggesting that the latter has a lower affinity for Pde6d. As the T26N mutation lowers the affinity of Rab28 for GTP, it can be inferred that GTP-binding promotes Rab28 association with Pde6d. To test this further, IPs were performed using just eGFP-Rab28^WT^, using treatment of the lysate with an excess of either GTPγS or GDP to force Rab28 into the respective conformations. Western blots with an anti-PDE6D antibody showed co-precipitation of Pde6d with all three Rab28 baits, however, in contrast to the MS data, the amount of Pde6d pulled down was the same for GTP*γ*S and GDP treatment and no treatment (**Figure 8D**). In summary, our data demonstrate that Rab28 interacts with phototransduction, membrane transport and mitochondrial proteins in the zebrafish larval eye.

## Discussion

### Loss of *rab28* leads to reduced cone OS shedding in zebrafish

Mutation of *RAB28* in humans is independently linked with cone-rod dystrophy in multiple pedigrees (Roosing et al. 2013; Riveiro-Álvarez et al. 2015; Lee et al. 2017). This form of retinal degeneration is characterised by initial cone death, followed by loss of rods. In agreement, a *rab28* knockout mouse displays cone-rod dystrophy, resulting from failure of cone outer segment (COS) phagocytosis (Ying et al. 2018). In the zebrafish knockout model described here, we also find perturbed COS shedding, resulting in a significant reduction in the number of phagosome-like structures positive for COS proteins within the RPE. We also find that overexpression of Rab28 GTP/GDP-preferring mutants does not significantly alter shedding. Our data confirms a conserved role for Rab28 in COS shedding. Given its broad conservation in vertebrates and eukaryotes generally, Rab28 likely acquired this function early in vertebrate evolution, possibly arising from a general role in the shedding of membrane (in the form of extracellular vesicles) from cilia (Jensen et al. 2016). The recent discovery that ectosome release is a conserved feature of cilia in many different cell-types and species provides an exciting opportunity to discover further regulators of OS shedding and phagocytosis. Indeed, one can speculate that photoreceptor outer segment phagocytosis is a specialised form of ciliary ectocytosis (Nachury & Mick 2019; Carter & Blacque 2019), as demonstrated for disc morphogenesis in mouse rods (Salinas et al. 2017).

### *rab28* mutant zebrafish have normal vision and retinal structure up to 12 mpf

In contrast to the mouse *rab28* knockout, *rab28* knockout zebrafish display decreased RPE phagosomes, but no retinal degeneration up to 12 mpf. One explanation is species differences in growth and regeneration. Unlike mammals, the zebrafish retina displays persistent neurogenesis throughout life, generating new cone photoreceptors at the periphery (Otteson & Hitchcock 2003). Retinal injury can also elicit a response from zebrafish Müller glia, which proliferate and re-differentiate to replace lost retinal cells (Yurco & Cameron 2005). These physiological differences may mask slow-onset thinning of the retina during degeneration.

Alternatively, genetic lesions which induce nonsense-mediated decay of mRNA were recently demonstrated to elicit a compensatory transcriptional response, whereby genes with similar functions are upregulated, masking the effect of the mutant gene (Rossi et al. 2015; El-Brolosy et al. 2019). This is particularly noted in zebrafish, where mutant phenotypes are often less severe or different from those of morpholino knockdown models (Kok et al. 2015), which do not display such compensation (Rossi et al. 2015). The absence of retinal degeneration in *rab28* knockout zebrafish may be the result of this compensatory transcriptional adaptation. Finally, there exists a high degree of redundancy in the zebrafish genome due to genome duplication thought to have occurred in the ancestor of teleost fish (Meyer & Van de Peer 2005). However, there is no indication for a *rab28* paralog in zebrafish. More globally, functional redundancy between different Rab family members may overcome loss of *rab28* (Pavlos & Jahn 2011; Blacque et al. 2018).

### Rab28 localisation in cones

Here, GFP-tagged Rab28 was highly enriched in the OS of zebrafish cones, regardless of whether it is in the GTP or GDP-bound state, suggesting all of its activity occurs within the OS. This agrees with previous findings that mouse Rab28 regulates shedding from cone OS tips (Ying et al. 2018). We also observed a striking pattern of localisation with COS, particularly those of short single (SS) cones, involving the concentration of Rab28 into discrete bands throughout the OS. This banding pattern was more extensive for the Rab28^T26N^ mutant, suggesting that GDP-bound or nucleotide free Rab28 is more efficiently targeted to these sites. As the average distance between cone OS lamellae (9-13 nm (Nilsson 1965)) is much too small to be resolved by fluorescence microscopy, these aggregations of Rab28 must be present on only a subset of lamellae within the OS. In rods and cones, a relatively consistent number of discs/lamellae are shed each time (Kocaoglu et al. 2016; Campbell & Jensen 2017). The precise reason for the *rab28* pattern in cones and whether this has a functional purpose are fascinating future research questions. One possibility is they mark sites of contact between the OS and the RPE. Rab28 may recruit effectors to these outer segment membranes which cooperate with other proteins in the RPE membrane to initiate outer segment shedding. Why this pattern is primarily observed in SS cones may be due to a need for higher disc turnover in UV COS, arising from their absorbance of highly phototoxic UV light. At the very least, our Rab28 localisation data indicates that there are differences in COS organisation between different classes of cone.

Discrete banding patterns within outer segments were previously reported, primarily in rods (Haeri et al. 2013; Hsu et al. 2015), though also in *Xenopus* cones for Prominin-1 (Han et al. 2012). Discrete patterns of localisation within COS are themselves surprising, given that proteins can freely diffuse both laterally and axially within them, via the ciliary facing membrane which connects lamellae (Young 1969; Liebman 1975). This is in contrast to ROS, where the separation of discs precludes diffusion of membrane proteins between them. Certain proteins, such as Rab28, must therefore be prevented from undergoing diffusion and tethered to particular membranes.

One implication from this is that photo-oxidatively damaged proteins are evenly distributed throughout COS, while they are restricted to the tip of ROS. Thus, OS renewal may be driven by entirely different mechanisms in cones and rods, potentially explaining why loss of *rab28* in mice appears to exclusively affect cone shedding and not that of rods.

### Rab28 interacts with the phototransduction machinery of zebrafish cones

Using an IP-MS approach, we identified novel interactants of Rab28 in the zebrafish eye. Of these, the most notable include components of the phototransduction cascade (opsins, Pde6c, Gucy2d, Gnb3b), vesicular trafficking proteins (Sv2a, Nsfa/b) and mitochondrial membrane proteins (Slc25a11, Atp5f1). We also validate a previously identified interaction with the prenyl-binding protein Pde6d. Interactions with phototransduction proteins may point to a role for Rab28 in the transport of these proteins, perhaps within the OS itself, given that Rab28 almost exclusively localises therein. The diversity of interactants either suggests roles for these proteins in photoreceptor OS, or roles for Rab28 outside the OS, such as in the inner segment (the location of photoreceptor mitochondria) or synapse.

There is a notably low degree of overlap between the interactants identified in our study and those in a previous study (Ying et al. 2018). There are several possible explanations for these discrepancies, first among them the characteristics of the species from which tissue was derived. The larval zebrafish retina is cone-dominant, consisting of ~92% cones (Zimmermann et al. 2018), in contrast to the rod-dominant bovine retina (Szél et al. 1996) used by Ying et al. Our dataset may therefore be more enriched for the cone-specific interactome of Rab28, as suggested by the prevalence of cone-specific proteins in our interactant list. Additionally, our experiments were performed with still-developing larval tissue. We show here that retinal development is accompanied by changes in Rab28 localisation within the COS and it is conceivable that this is accompanied by changes to the Rab28 interactome. Our data also offers insight into the effect of nucleotide binding on Rab28 interactions, as a small number of interactants only displayed significant interaction with one of the nucleotide binding mutants. For example, Gnb3b was significantly enriched by the Q72L (GTP-preferring) bait only, while Gucy2d and Erlin-1/2 were enriched for the T26N (GDP-preferring) mutant. Gnb3b may be a direct effector of Rab28 in its active state, while Erlin-1/2 may be a guanine nucleotide exchange factor (GEF), which promotes the exchange of GDP for GTP (Lee et al. 2009).

## Acknowledgements

We thank Ross F. Collery for providing the reagents for the Tol2 kit and David Hyde and Susan Brockerhoff for generously providing anti-zebrafish opsin and anti-cone transducin alpha antibodies, respectively. We also thank the UCD Conway Institute Imaging and Proteomics core facilities for imaging and mass spectrometry support and services, respectively. Funding for this research was provided by Irish Research Council grant GOIPG/2014/683. DM is funded by Science Foundation Ireland 15_CDA_3495.

---

**Supplementary figure 1.**
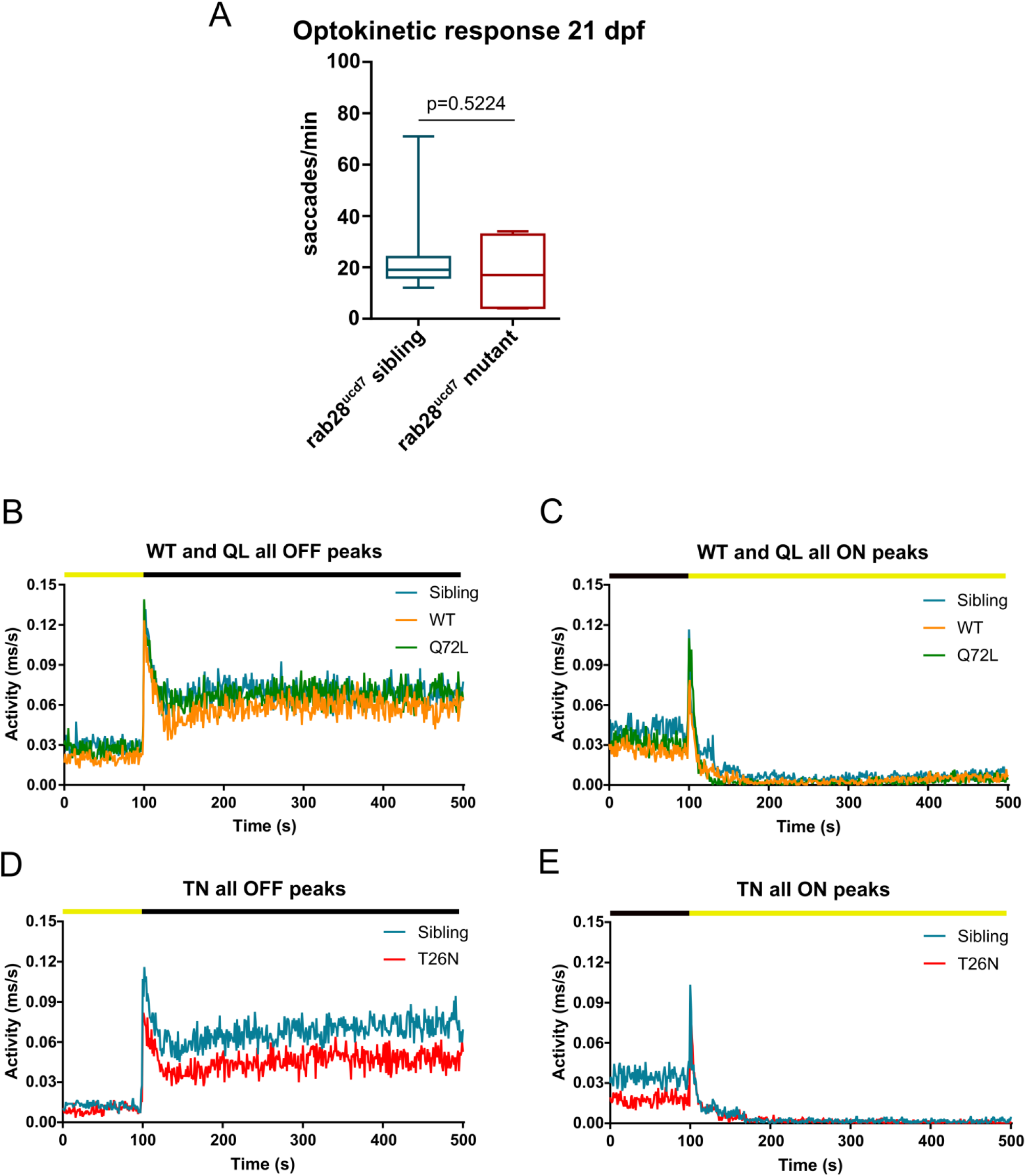
(related to figures 4 and 5). 21 dpf OKRs and transgenic larvae VMR peak activity traces. **(A)** Box and whisker plot of optokinetic response (OKR) assay of 21 dpf *rab28* mutants, sibling controls and eGFP-Rab28 transgenic zebrafish. Data are from 4 *rab28* mutants and 11 controls. **(B-E)** Activity traces of 5 dpf transgenic and sibling larval activity during the VMR assays, around the times of peak OFF and ON activity. Black and yellow bars indicate dark and light conditions, respectively.

**Supplementary figure 2.**
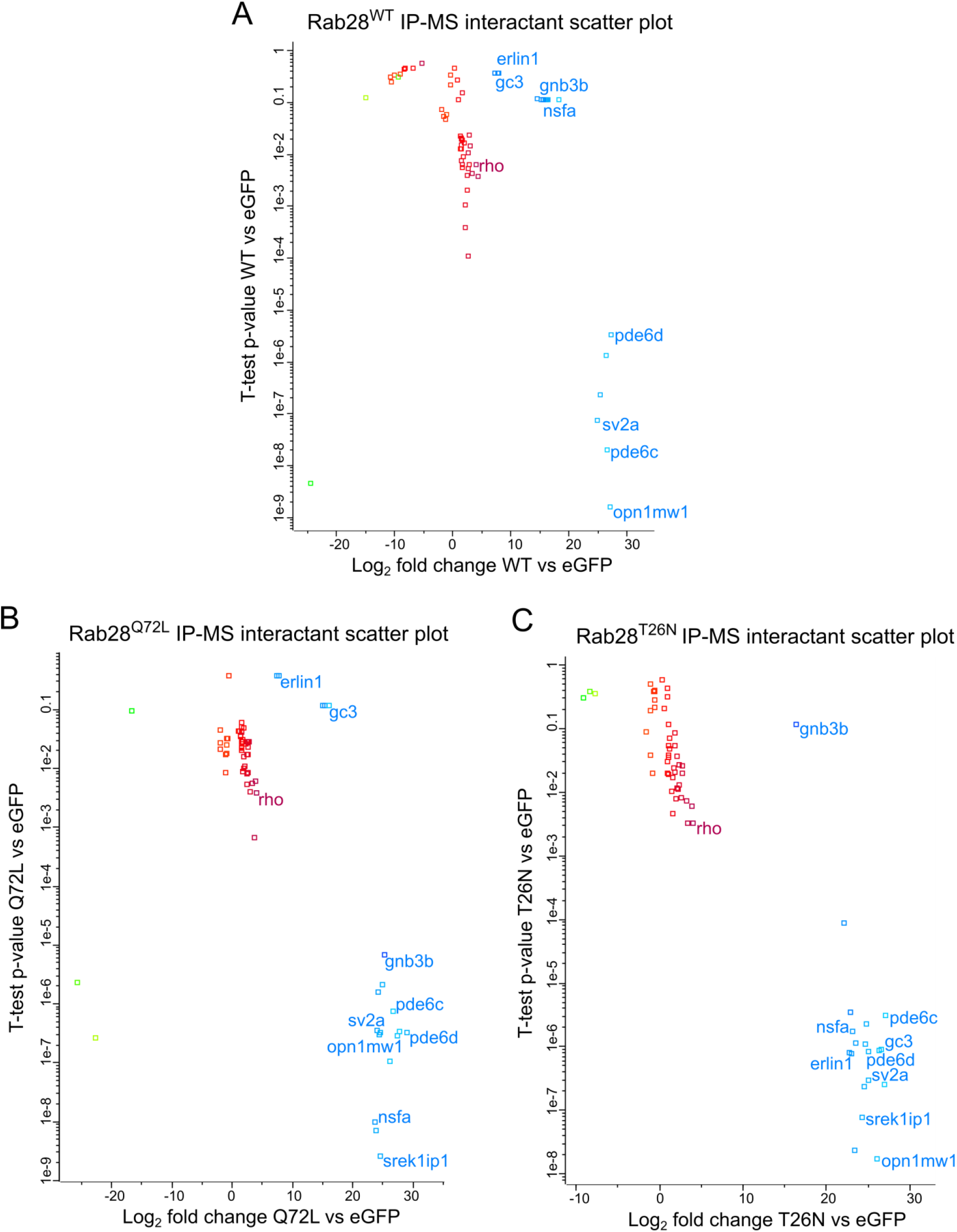
(related to figure 8). Scatter plots of MS data for identified Rab28 interactants. **(A-C)** Scatter plots showing the p-value vs log2 fold change of proteins identified by Rab28 IP-MS. Proteins group into two clusters, one with high fold change and significance and one with a low fold change and significance.

